# Revolutionizing large-scale DNA synthesis with microchip-based massive in parallel synthesis system

**DOI:** 10.1101/2024.10.30.619547

**Authors:** Xiandi Zhang, Xianger Jiang, Yun Wang, Qinzhen Chen, Ruihong Zhang, Hao Jiang, Hu Zhang, Antoni Beltran, Weiya Yang, Chenglong Liang, Ning Chen, Yun Huang, Guqiao Ding, Chengwang Xie, Nanfeng Gao, Kaijing Zheng, Juntao Liu, Wei Xu, Jinlei Huang, Dong Cai, Lihao Zhu, Songjin Mo, Mengzhe Shen, Wenwei Zhang, Ben Lehner, Ming Ni, Jian Wang, Xun Xu, Yue Shen

**Author notes:** Correspondence (Y.S.). These authors contributed equally (Xiandi Zhang, Xianger Jiang).

## Abstract

DNA synthesis serves as the fundamental enabling technology of engineering biology, aiming to provide DNA molecules of designed composition, length, and complexity at scale and low cost. Current high-throughput DNA synthesis technologies rely on intricate chip manufacturing and microfluidic systems to provide large-scale synthetic oligonucleotides, at the expense of low concentration and limited compatibility in the processing of longer DNA constructs assembly. Here, we report a microchip-based massive in parallel synthesis (mMPS), pioneering an “identification-sorting-synthesis-recycling” iteration mechanism to microchips for high throughput DNA synthesis. In comparison to microarray-based methods, we demonstrate that our method can increase the DNA product concentration by 4 magnitudes (to picomole-scale per sequence) and greatly simplifies the downstream processes for large-scale gene synthesis construction. By the construction of 1.97 million-diversity variant libraries that cover 1,254 human protein domains, we demonstrated the uniformity of the constructed variant libraries using mMPS-derived oligos is greatly improved, with amino acid distribution highly consistent as designed. In addition, by synthesizing 285 1kb-to-3kb genes with varying degrees of sequence complexity from previously reported strains A501 and 3DAC, potential ancestor of early archaea and bacteria, our result shows that the overall gene assembly success rate using mMPS-derived oligos is increased by 10-fold in comparison to other methods. Our mMPS technology holds the potential to close the gap between the quality and cost of writing DNA in increasing demand across many sectors of research and industrial activities.

## Introduction

Our growing capability to engineer biological systems is set to radically transform the ways we understand biology. For example, artificial intelligence (AI) algorithms have shown remarkable success in predicting protein folding and interactions^1^, the application of engineering biology tools, in conjunction with AI and data-driven computational techniques, enables researchers to explore vast sequence space with unprecedented accuracy, allowing for the rapid identification of protein sequences and structures that link to specific functions. Today, reconstructions of complete bacterial and eukaryotic genomes^2-6^ show the potential of assessing the functional diversity and evolutionary path of bioresources or creating novel organisms tailored for diverse applications. In addition, the convergence of biotechnology and information technology catalyzed the development of DNA-based data storage, in which large-scale DNA sequencing and synthesis are used for information reading and writing processes^7^. Consequently, the demand for synthetic DNA is increasing in both fundamental and applied biological research.

Especially nowadays, a vast majority of biological research and engineering biology studies rely on synthetic DNA, which is not limited to oligonucleotides (oligos), but also longer constructs like genes or even entire chromosomes^8^. There is a shift towards enzymatic DNA synthesis, which holds the potential to synthesize longer oligos with greener solutions^9^. However, the current throughput and cost of this emerging technology limits its wide applications with large-scale demand. Massively parallel oligo synthesis on an arrayed chip surface employing solid-phase phosphoramidite chemistry (also known as DNA microarray) remains the most popular method^10^, leading to orders of magnitude reduction of the cost than the widely commercialized column-based method^11^. The spatially localized DNA synthesis on the arrayed chip surface can be achieved by several different strategies, such as light-based chemistries, ink-jet printing, and semiconductor-based electrochemistry^12-15^. The fact is that DNA microarray is originally developed for the purpose of high throughput genome-wide gene expression profiling and target capturing for next-generation sequencing^12^. Thus, all the synthesized oligos on the arrayed surface are cleaved and harvested as a ‘oligo pool’, in which the number of oligos can be very large.

Existing technologies lead to two major challenges in using the microarray-based oligo pools to build longer constructs. First, the concentration of oligos cleaved from the solid support typically falls in the picomolar range, with individual sequence yields often at the femtomole (fmol) scale^16^. Such low yields are insufficient for direct application in gene synthesis, underscoring the need for improved production strategies, (see section “mMPS system improves the success rate of high throughput gene assembly”). Therefore, PCR amplification is required to increase the concentration of oligos prior to assembly. Considering the error rates for oligo pools are usually higher than column-based strategy, introducing the PCR step would further amplify the errors in the resultant raw materials for gene synthesis. Consequently, enzymatic error correction of the assembly products is necessary and brings additional cost to the whole process. Second, at greater pool complexities, constructing any individual gene in parallel becomes very challenging mostly owing to spurious cross-hybridization of oligos in a pool. To address this issue, several approaches have been developed. One is the introduction of predesigned barcodes adjacent to the designed oligos^17^, which allowed PCR amplification of oligos participating in only a particular assembly. However, that would result in substantial more synthesis efforts in comparison with the actual need. In addition, the design of barcodes will be even more challenging if the gene sequences are similar to one another. Another approach is customization of the arrayed surface with physically separated microwells for the synthesis of oligo subsets^18^, but at the expense of relying on intricate chip manufacturing. Either way will lead to increased cost to make longer construct and is difficult to further scale up.

To address the forementioned challenges, we report a new strategy called microchip-based massive in parallel synthesis (mMPS), which applies millimeter-size microchips for high throughput DNA synthesis through an “identification-sorting-synthesis-recycling” iteration mechanism. The construction of variant libraries covering 1,254 selected human protein domains and 285 1kb-to-3kb genes with varying degrees of sequence complexity from previously reported potential ancestor of early archaea and bacteria using mMPS derived oligos validated the feasibility of this strategy. In comparison with microarray-based approach, the concentration of mMPS-derived oligo products is significantly increased by at least four orders of magnitude, reaching pmol-scale. And the gene synthesis steps are also greatly simplified into only two steps, leading to a 10-fold success rate of assembly. With advantages at scalability and flexibility of throughput, our mMPS technology holds the potential to close the gap between reading and writing DNA, facilitating the test of generated data-driven hypotheses experimentally at unprecedented scale and low cost.

## Results

### Barcoded microchip enables high-throughput DNA synthesis with a sortable process

Among current microarray technologies, high throughput is accomplished by creating arrays with micrometer or sub-micrometer reaction spots on glass or silicon wafers^14, 19, 20^. However, increasing the density of these reaction spots, while enhancing throughput in oligo synthesis, typically leads to a reduction in yield as the surface area is decreased. Furthermore, the pursuit of ultra-high throughput synthesis not only compromises the yield of individual sequences but also necessitates greater precision in array fabrication and a more intricate liquid handling system. To overcome these constraints and to offer flexibility in throughput, our system is developed by using customized millimeter-size microchip to serve as the solid support for oligo synthesis in a traceable manner, allowing to add bases in series from 3′ to 5′ one at a time on the customized mMPS microchip as solid support over a series of ‘Identification-Sorting-Synthesis-Recycling’ cycle by employing the standard phosphoramidite chemistry^21^ (**Figures 1a&1b**). Comparing with other arrayed chip-based techniques that generate oligo pool on one chip (multiple oligo sequences vs. one chip), the most different point of mMPS system is that one microchip generates one oligo (one oligo sequence vs. one microchip). The system’s throughput is directly subject to the quantity of mMPS chips employed simultaneously for each run. And the current upper limit of throughput for each run is at one million (the encoding capacity of the QR code).

**Figure 1.**
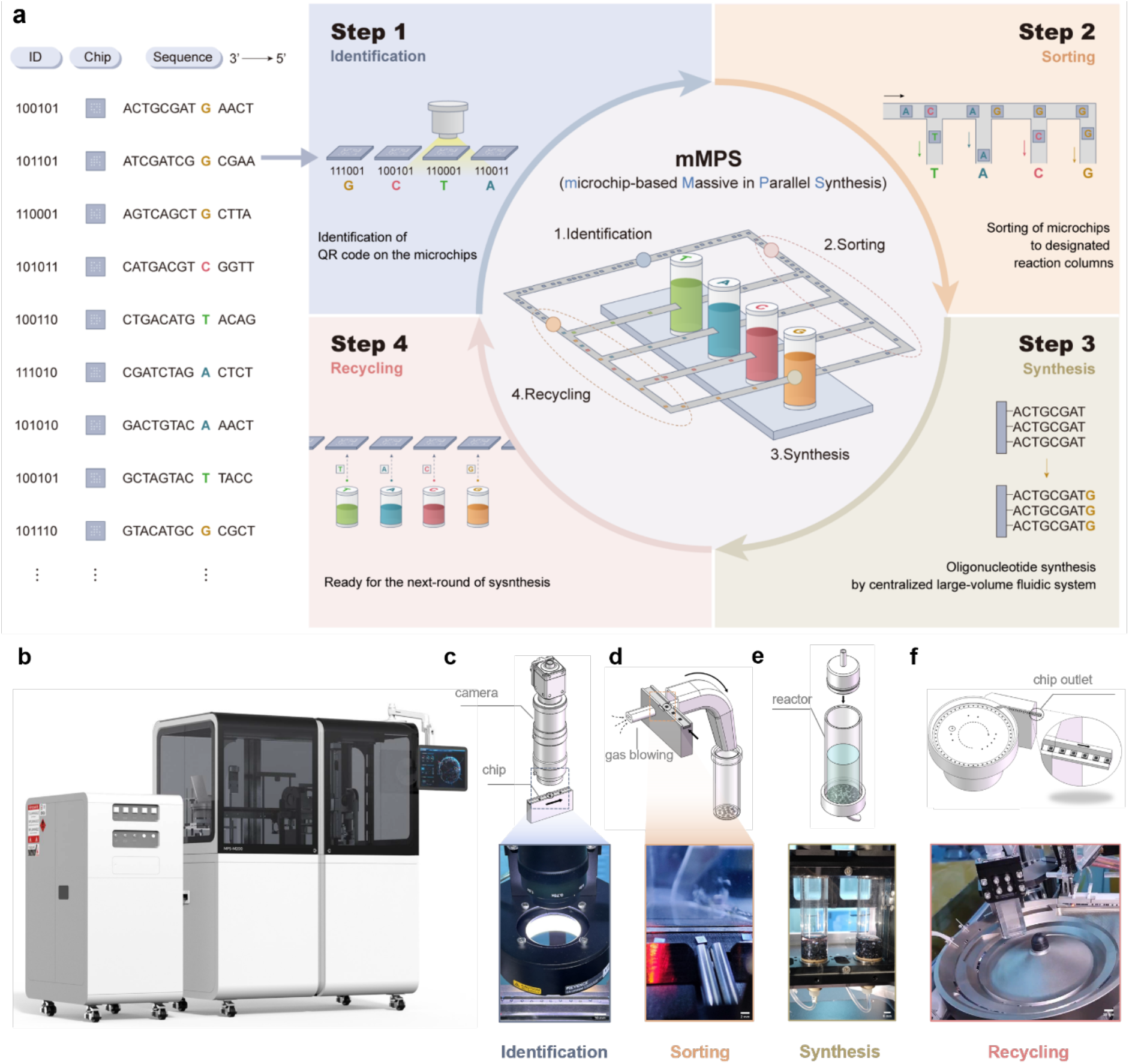
The barcoded microchip based Massive in Parallel Synthesis (mMPS) method allows high throughput DNA synthesis in a sortable approach. Schematic illustration of the (a) operational principle of the mMPS method and (b) integrated mMPS system. Schematic illustration and digital image of (c) identification, (d) sorting, (e) synthesis, and (f) recycling module.

For identification, high-speed camera is used to detect each mMPS microchip marked with a unique QR code on its surface (**Figure 1c**). Then as a response, the detected microchip is sorted to the corresponding synthesis reactors (sealed cylindrical columns) to complete the synthesis step (**Figure 1d**). The synthesis process of each type of base is completed in its dedicated columns, with inner gas protection employed to exclude the influence from moisture (**Figure 1e**). Additionally, the fluidic system adjusts reagent volumes dynamically based on the quantity of introduced microchips, minimizing reagent consumption waste. After nucleotide addition, recycling module transfers microchips back for the next cycle (**Figure 1f**). Notably, this design of dedicated synthesis reactor for each type of nucleotide in mMPS not only prevents cross-contamination among nucleotides but also allows for the optimization of reaction conditions, *e.g*. deprotection time, base concentration, and coupling duration, *etc*., tailored to each nucleotide. Consequently, the mMPS system is able to achieve a customizable reaction process designed to ensure high-quality performance and flexible throughput across diverse sequences and applications.

### Design strategy in mMPS microchip and performance evaluation in oligonucleotide synthesis

In comparison to conventional arrayed chips in high-throughput DNA synthesis, design strategies of solid support are limited to manufacturing process, often neglecting the substantial influence of the chip’s surface chemistry and physics^22, 23^. In solid-liquid chemistry, however, surface properties of solid support largely affect the reaction performance by affecting the interactive dynamics at the interface^24, 25^. As illustrated in **Figure 2a**, the microchip’s design is crucial for determining the throughput, stepwise efficiency, and yield of each synthesized sequence. While several methods have been developed to enhance stepwise efficiency and throughput^19, 26^, strategies to increase yield are underexplored. The yield of individual sequences is determined by the number of active sites available for nucleotide elongation, which is inherently dependent on the microchip’s design.

**Figure 2.**
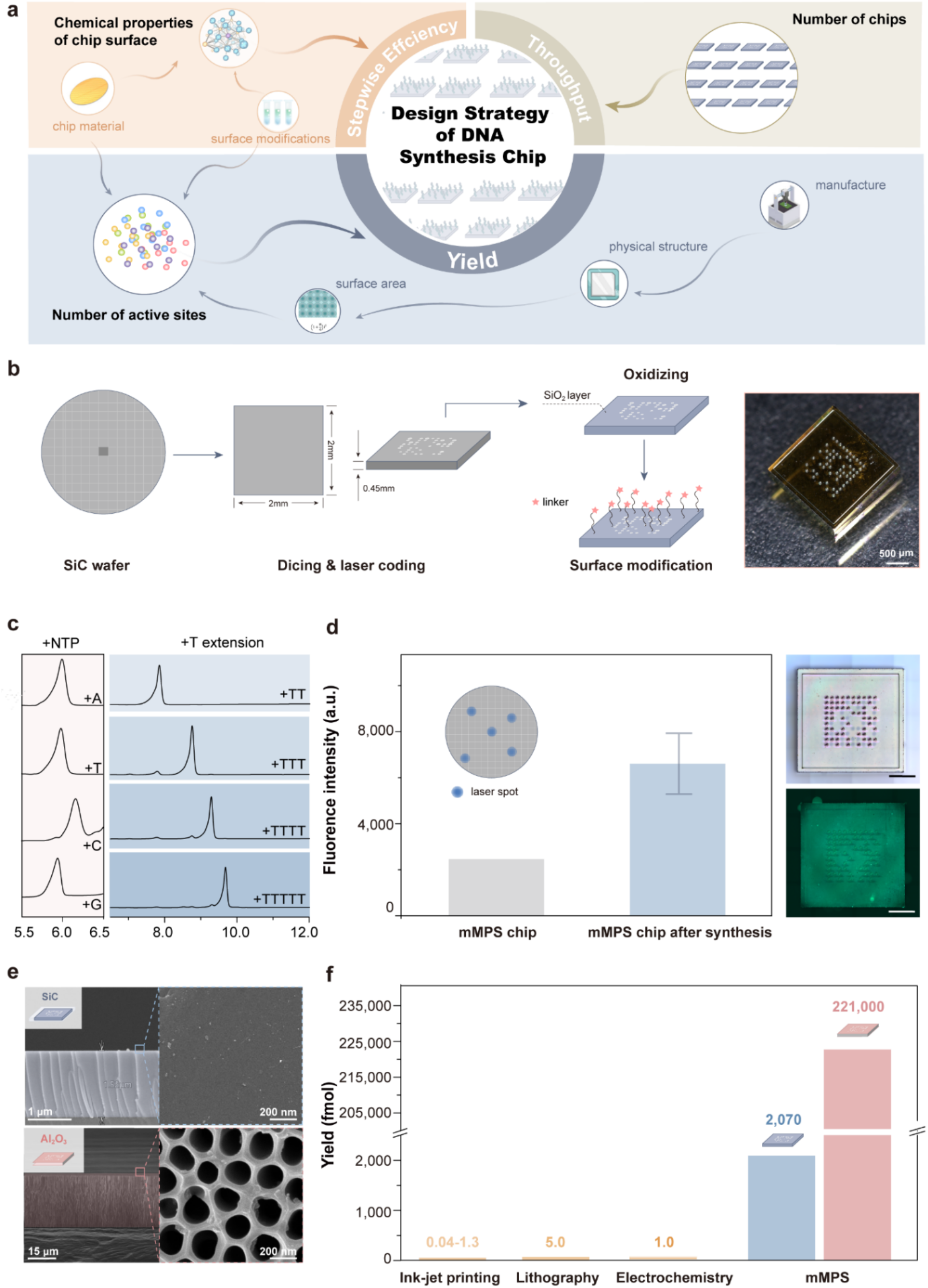
The design and manufactures of mMPS microchip generate oligo product at pmol to nmol scale. (a) Schematic illustration of chip design strategy. (b) Manufacturing process (left) and depth camera image of microchip (right). (c) Feasibility evaluation for the synthesis of distinct nucleotide bases (left) and the extension by the addition of thymine (right) on microchip using HPLC analysis. (d) PL spectra (left) and fluorescence images (right) of microchip post-oligo synthesis. (e) Schematic and high-resolution SEM image of porous microchip, (f) Yield comparison of synthesized oligos with other existing methods.

In light of these considerations, we have meticulously engineered the physical structure and chemical properties of the mMPS microchip, aiming to improve the produced oligo yield. We selected a silicon carbide (SiC) wafer as the substrate for the microchip due to its superior stability and amenability to manufacturing processes^27^. The SiC wafer was functionalized into microchips through a series of processes, including laser dicing, wet-oxidation, and chemical modification (**Figure 2b**). Characterization of the microchip, conducted using X-ray diffraction (XRD), scanning electron microscopy (SEM), and ellipsometry, revealed its composition to be SiC with a uniformly formed amorphous silicon dioxide (SiO_2_) layer of 1021.6 nm in thickness on the surface. This configuration results in a homogeneous SiO_2_@SiC core-shell structure within the microchip, which is integral to its enhanced performance in high-throughput DNA synthesis (**Figures S1, S2** and **Table S1**). To evaluate the feasibility and performance of oligo synthesis with modified microchips, a series of demonstrations were conducted *via* mMPS system. We commenced by testing the mMPS microchip’s capacity to execute a single cycle of DNA synthesis for each deoxyribonucleotide triphosphate (dNTP). High-performance liquid chromatography (HPLC) analysis confirmed the successful synthesis of the desired products (**Figure 2c**, left). Subsequently, we take polyT extensions to assess the mMPS system’s capability in sequence synthesis. The HPLC spectra clearly demonstrated the incremental synthesis progression from dinucleotide (T2) to pentanucleotide (T5) stages (**Figure 2c**, right), and further extended to decanucleotide (T10) and trinucleotide (T30) sequences (**Figure S3**), achieving the targeted sequence lengths with precision.

Post-synthesis quality assessment of the products on the mMPS microchips was conducted using SYBR Gold staining, followed by photoluminescence (PL) spectroscopy and fluorescence microscopy analysis. **Figures 2d** and **S4** depict uniform excitation peaks at approximately 530 nm, attributable to the SYBR Gold fluorescence signal from the mMPS microchips post polyT synthesis. In contrast, the planar chip exhibits a peak at approximately 460 nm with diminished intensity, indicative of the intrinsic SiO_2_ surface excitation. Further fluorescence imaging visualizes homogeneous signal distribution, indicative of the uniform ssDNA growth observed (**Figure 2d**, right). Notably, the calculated yield of the target sequence, as derived from HPLC spectra, is 2,070 ± 157 fmol with a stepwise efficiency of 99.3%, thereby validating the design strategy employed in the mMPS microchip. To further explore the potential of increasing yield, we engineered a three-dimensional porous microchip based on Al_2_O_3_. This microchip features an optimized thickness of 35 μm and a pore diameter of 218 ± 33 nm (**Figures 2e and S5**). With this design, the synthesis of a T30 sequence on the porous mMPS chip yielded 221,000 ± 415 fmol of oligo production. Compared to existing DNA synthesis methods, the mMPS system’s specific microchip yields are 4 to 6 orders of magnitude higher than most current techniques (**Figures 2f** and **Table S2**), offering the opportunity to revolutionize downstream applications by enabling direct application without the necessity for PCR amplification.

### mMPS system improves the success rate of high throughput gene assembly

High-throughput gene synthesis is fundamental to advancements in synthetic biology and biotechnology. Polymerase Chain Assembly (PCA) is particularly prominent for synthesing gene length ranging from a few hundread base pairs to several kilobase pairs due to its simplicity and high assembly success rate^28, 29^. Previous efforts have demonstrated the feasiability of using PCA to assembly genes with arrayed chip derived oligos^18^. Generally, the overall success rate of PCA is influenced by several key factors, including proper concentration, number, and length of the oligos.

Since arrayed chip-derived oligos are produced as a mixture pool and also at very low yields. Consequently, introducing multiple pairs of barcode primers is required in order to increase the oligo concentration and also reduce the complexity of assembly caused by spurious cross-hybridization of oligos (**Figure 3a**). In comparison, mMPS system can avoid the introduction of barcode primers since each chip carrying the generated oligos required for one gene can be physically sorted, cleaved to obtain the oligo premix for direct assembly, leading to a significantly simplified assembly workflow (**Figure 3a**). In addition, by serial dilution of column-based method derived oligos, we found that the at least 200 fmol of each oligo in a 20 μL PCA reaction is required to ganrantee successful assembly of genes (**Figure S6**). Considering a certein loss of oligo product during the sample preparation for assembly, the yield of mMPS derived oligo yield perfectly matches the requirement for gene assembly, reducing the risk of unintended mutation introduced or error accumulation during PCR amplificaiton step.

**Figure 3.**
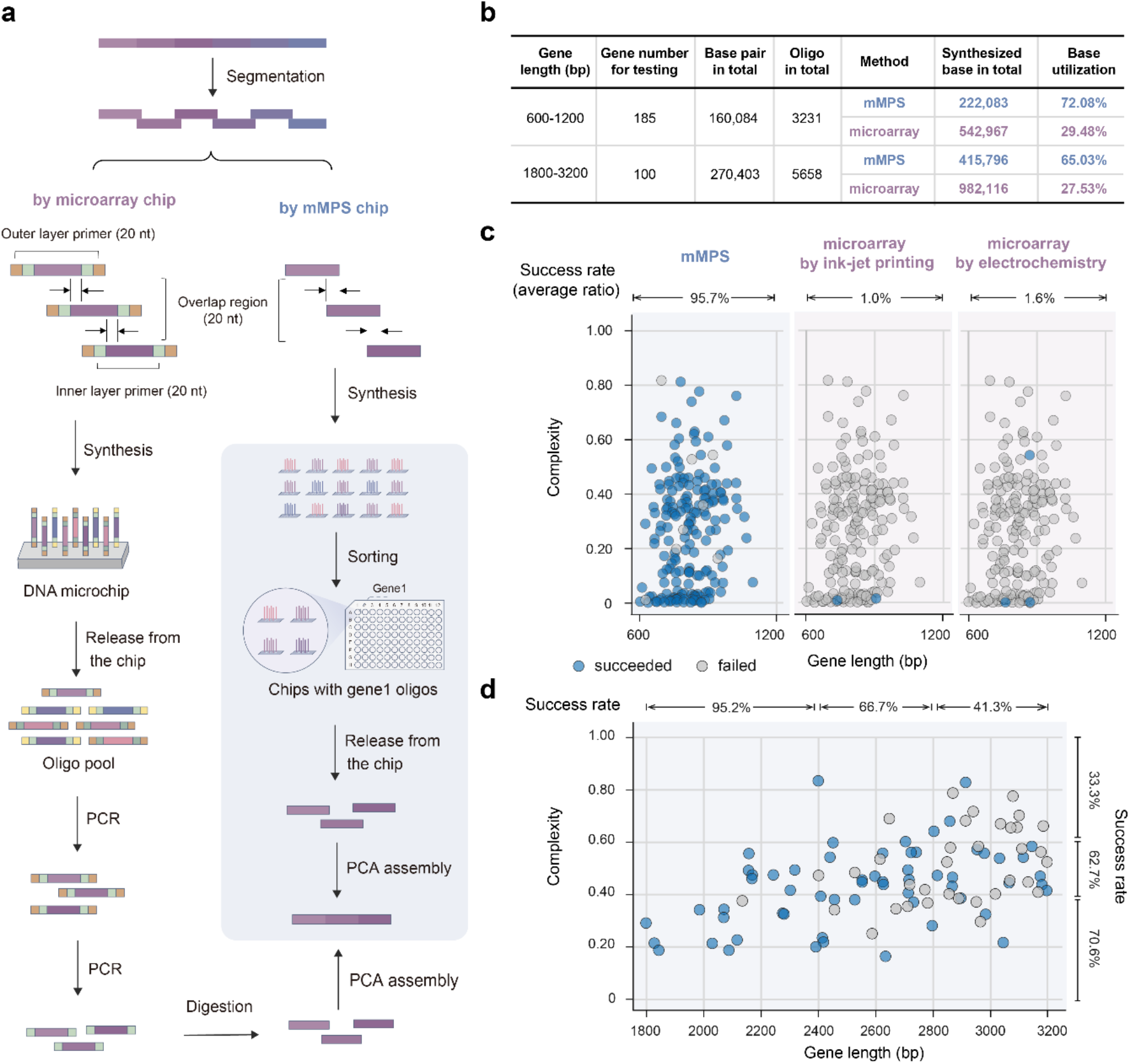
Comparison of Gene Synthesis using mMPS system and two representative high-throughput arrayed chip DNA Synthesis methods. (a) Schematic illustrating gene assembly using conventional arrayed chip derived oligos and mMPS oligos. (b) Statistics of oligo design for 600-to-3200 bp genes. (c) Average success rate for ∼1 kb genes assembly using mMPS derived oligos and two representative arrayed chip derived oligos. (d) Average success rate for for ∼2-3 kb genes assembly using mMPS derived oligos.

Next we selected 285 genes ranging from 600 bp to 3,200 bp from two previously reported strains A501^30^ and 3DAC^31^, potential ancestor of early archaea and bacteria, to test whether the performance fits to our hypothesis in comparison with two representative arrayed chip based DNA synthesis methods. In addtion, these genes also represent varying levels of sequence complexity, which takes repeats, secondary structures, GC% and Tm extremes into consideration^32^. The gene sequences were first segmented into in total 8,889 of 41-79 nt oligos for assembly (**Figure 3b**). In additon, since array chip based DNA synthesis methods require post-amplication and restriction enzyme digestion, two pairs of 20-nt barcode primer sequence and restriction digestion sites were introduced at the 3’ and 5’ ends of generated oligos. That leads to a drastic reduction of synthetic base utilization rate, which refers to the total amount of bases for synthesis divided by the actual total amount of bases been synthesized. For shorter genes (from 600 to 1,200 bp) and longer genes (from 1,800 to 3,200 bp), the correspondingly synthetic base utilization rate of arrayed chip based approach is only at 29.48% and 27.53% respectively, while that of our mMPS based approach increases to 72.08% and 65.03% respectively (**Figure 3b**), suggesting our method can greatly improve the economic benefit of gene synthesis manufacture process.

Then we synthesized the 3,231 designed oligos for all 600-to-1,200 bp genes by both mMPS system and two representative arrayed chip DNA synthesis technologies (by ink-jet printing and electrochemistry). Following the corresponding workflow of assembly, we verified and compared the overall success rate of gene assembly among the three selected technoloies. While we demonstrated 177 out of 185 genes can be successfully assembly using the mMPS derived oligos, leading to a success rate at 95.7% (**Figure 3c, S7**). It is astonishing to see that only very few genes can be assembled, showing the right size on the gel, by ink-jet printing approach or electrochemistry approach (**Figure S8, Figure S9**), with their corresponding success rates both less than 2% (**Figure 3c**). Our result demonstrated that the assembly efficiency can be greatly influenced by the complexity of assembling workflow. Then we want to further explore whether our mMPS system can assemble longer genes through one-pot PCA assembly. We synthesized the rest 5,658 generated oligos for the 100 genes at 1,800-to-3,200 bp length by our mMPS system. We observed a clear downtrend of assembly success rate as the gene length gets longer and sequence complexity gets increased(**Figure 3d, S10**). However, the overall success rate of mMPS based approach far superior to the other two arrayed chip based methods. Our result demonstrated that our mMPS system can facilitate gene assembly process in a rapid, cost-effective manner while achieving a high success rate, holding the great potential to support the next milestone achievement of genome scale writing efforts.

### Construction of human protein domains variant libraries at ∼2 million diversity with mMPS system

With advances in AI and generated big data, the demand for variant libraries to facilitate protein functional studies and genetic engineering has surged. Traditional techniques such as site-directed mutagenesis (SDM)^33^, error-prone PCR^34^, and Tn5-mediated transgenesis^35^, are inherently limited in their ability to introduce only a limited number of mutations simultaneously^35^. This constraint hinders our capacity to comprehensively investigate genotype-phenotype relationships. The development of massively parallel DNA synthesis technology has enabled deep mutational scanning (DMS)^36^, such as NNN/NNK-based libraries^37^, by which extensive variants at designated site can be introduced for investigation. Generally, a variant library with high coverage, uniformity and fidelity is crucial for generating high quality data to deepen our understanding of protein structure and evolutionary dynamics, thereby improving our ability to predict protein interactions and functional impacts. Previously, variant libraries generated using array chip-based methods encountered two main issues. First, the low yield at the fmol scale of generated oligos was inadequate for multiple attempt experimental applications, then batch-to-batch variability might further affect the reproducibility of results. Second, the uniformity of the introduced variants (e.g., N = 25% A, 25% T, 25% C, 25% G) is uneven, which might introduce bias in subsequent experiments and potentially lead to incorrect conclusions.

To test whether our mMPS system can address these issues, we chose to build a NNK variant libraries used in another study, which aiming at performing the first large-scale site saturated mutagenesis to human protein domains to investigate the consequence of human missense variants to protein stability and build the reference dataset for clinical variant interpretation^38^. Specifically, 1,254 human protein domains were selected and designed to have NNK-type codon mutational scanning (i.e. every amino acid mutated to all other 19 amino acid at every position in each domain), leading to nine libraries with length ranging from 101 to 335 bases and a total library diversity approaching two million (**Figure 4a,4b**). The degenerate bases (N/K) mixed the deoxynucleotide with an optimized ratio, ensuring an equal representation of amino acids at mutation sites and enabling an unbiased assay for mutational effects. For the degenerate base synthesis, two additional synthesis reactor columns were used during the synthesis process. The constructed variant libraries by mMPS system were first cloned into plasmid and transformed into yeast cells with three replica repeats for sequencing. The abundance measurements across all libraries shows high reproducibility, with a median Pearson correlation coefficient of r=0.97 between replicates (**Figures S11**).

**Figure 4.**
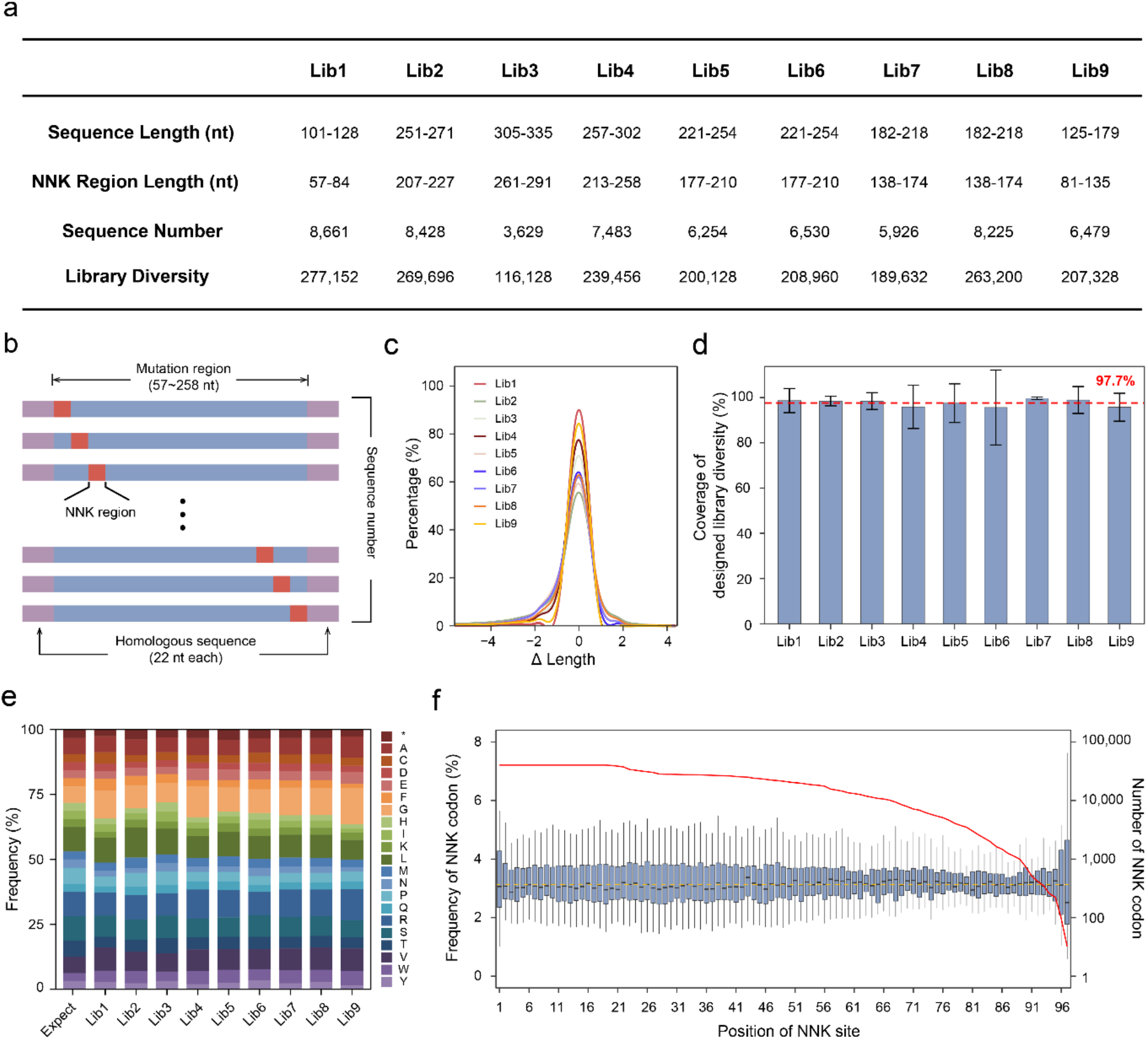
Performance of mMPS-generated deep mutational scanning libraries. (a) Summary of library construction in nine libraries (Lib1 to Lib9). (b) Schematic of deep mutational scanning library design. (c) Analysis of sequence length deviation (ΔLength) from the expected length across libraries. (d) Coverage analysis across each library. (e) Amino acid distribution analysis for each library, with theoretical expectations displayed on the left. The * symbol denotes stop codons. (f) Distribution and coverage of NNK codon across targeted sites of all libraries. The blue box indicates the frequency of NNK codons across targeted mutation sites, the line inside the box marks the median frequency. The red line shows the total number of NNK codons observed per position for the nine libraries. The brown dotted line indicated the expect frequency of NNK codon.

Then we analyzed the sequence length distributions across all libraries (**Figure 4c**). The deviation from expected length (ΔLength) demonstrated a tightly centered distribution around zero, with a substantial proportion of sequences (approximately 70%) displaying no length deviation. Notably, Lib1 demonstrated an exceptional performance, achieving up to 90% with full-length sequence. The high fidelity in length maintenance across libraries effectively avoids frameshift mutations during protein translation, thereby enhancing the effective utilization rate of the mutant libraries. Next, we assessed the mutant coverage by comparing to the designed library diversity. Across all libraries, the coverage ranged from 95.5% to 99.7%, with an average coverage rate at 97.7% ± 1.6% (**Figure 4d**). In addition, to evaluate the uniformity of library, we analyzed the amino acid distribution in the domain regions, which confirmed a nearly consistent frequency profile aligned with theoretical expectations. The average coefficient of variation (CV) in the fold change of observed frequency compared to expectation was 0.25. Our results indicated that all expected variants were represented at the desired ratios without bias (**Figure 4e, Figure S12**), which is significantly better than the same type of variant libraries synthesized through two other commercialized vendors (**Figure S13**). To further assess the distribution of NNK codons across targeted mutation sites, we calculated the frequency of NNK codons at each position within the mutagenized region. Despite variations in sequence lengths across the nine libraries (**Figure 4a**), which led to fewer NNK codons at terminal positions, the majority of sites demonstrated a uniform distribution, reflecting consistent mutational coverage (**Figure 4f**). These results underscore the high coverage, uniformity, and fidelity of libraries constructed using the mMPS system, which offered strong support the downstream investigation efforts that successfully reported the effects of more than 500,000 missense variants on the stability of over 500 distinct human protein domains^38^.

## Discussions

The field of engineering biology is rapidly evolving, increasing the demand for a high-throughput, cost-effective DNA synthesis method. However, the current approaches to high-throughput oligonucleotide synthesis utilize similar routes, lacking of breakthrough on synthesis performance for a long time, suggesting a slow pace of innovation. In addition, the material structure involved in arrayed chip-based DNA synthesis is typically uniform, and the preparation methods are often homogeneous. Despite being a critical component in oligo synthesis technology, few studies address the rational design of novel synthesis chips, with limited scientific explorations of the principles behind. We innovatively introduce a substrate for the synthesis chip based on a non-silica material and engineer its surface structure to significantly enhance the production yield of individual oligo. We successfully demonstrated that our mMPS system dramatically increases DNA product concentration by 4 to 6 orders of magnitude to the pmol to nmol scale per sequence, together with its unique sorting characteristic by the employment of QR code, simplifying the processes and improving the quality for large-scale longer construct synthesis. This indicates that a rational design of chip physical and chemical properties could greatly improve synthesis performance. Besides, Enzymatic synthesis methods are emerging but have yet to achieve high-throughput capabilities. Integrating the mMPS system with enzymatic techniques holds promise to produce longer oligonucleotides as well as presents a more sustainable alternative, mitigating the environmental impact associated with the use of organic solvents and hazardous chemicals prevalent in conventional DNA synthesis methods. The error rates observed in mMPS fall within acceptable limits for applications such as variant library construction and high-throughput gene synthesis, demonstrating notable improvement over previously reported values without error correction. This outcome suggests a compelling potential for further enhancement through the integration of an error-correction step in subsequent processes. We anticipate that incorporating error correction could reduce synthesis error rates even further, thereby broadening and deepening the applicability of the mMPS DNA synthesis system across a wider range of research and industrial applications.

## Online Methods

### Chip Preparation on mMPS chip and mMPS_porous chip

The mMPS chip was fabricated by Hopengine Technology, Inc. from 6-inch SiC wafer. Typically, the wafer was initially sectioned into squares measuring 2×2×0.45 mm, followed by laser perforation on both faces of each square. Subsequently, each square was marked with a distinct QR code *via* laser perforation on both sides. The perforated substrates underwent wet-oxidation at 1000 °C for a duration of 60 h. Post-oxidation, the square substrates were sonicated with deionized water and acetone for three times and dried in the oven under 75 °C for 15 min, namely mMPS chips.

The mMPS porous chips were developed from alumina plate with a thickness of 0.40 mm. The alumina plate is firstly labeled with QR code in an array of 2×2 mm squares. Post labelling, electrochemical method was employed to develop Al_2_O_3_ porous structure on both surface of alumina plate^39^. Laser perforation was used to dice the plate into 2×2 mm squares with the pre-marked code reserved. Then the aluminate-based squares were sonicated with deionized water and acetone for three times and dried in the oven under 75 °C for 15 min, namely mMPS_porous chips.

### Chemical Modification of mMPS chips and mMPS_porous chips

Chemical modification of mMPS chip was proceeded with silanization and linker connection. In a typical procedure, 30,000 chips were sonicated in 150 mL of prepared silane solution (0.9 vol.% APTES/ 0.9 vol.% PTES/acetone) for 120 min and washed with acetone for 5 times, followed by drying in the oven under 75 °C for 15 min. After silanization, 2.42 g of universal linker and 2.84 g HATU were dissolved in ACN, following adding 7.1 mL of DIPEA and mixed as linker solution. The silane microchips were sonicated with linker solution for 120 min and washed with ACN and acetone three times, respectively. Treated chips were dried in the oven under 75 °C for 15 min. Chemical modification of mMPS_porous chips is slightly different where the silanization of 1,000 chips is proceed at 90 °C for 3 h in 100 mL of prepared silanization reagent (0.5 vol.% APTES/ 0.5 vol.% PTES/ toluene).

### Oligo synthesis on mMPS system

Before synthesis, the file containing sequence information that is about to synthesis was imported into the mMPS operation system. Oligo synthesis through mMPS is divided into four steps and processed automatically, depicted as follows.

Step 1: Identification:

A certain amount of mMPS chips were aligned by a vibratory bowl. Within the bowl, a carefully calibrated vibrational action reorganizes the chips, ensuring they are arranged in a neat, single-file formation for a systematic exit through the outlet. Subsequently, the aligned chips are transported by a rubber belt, which maintains a consistent rate of transfer, allowing the camera to reliably capture and identify each chip.

Step 2: Sorting

Once the QR code has been recorded by the identification module, chips would be automatically linked with a pre-determined sequence information based on the uploaded synthesis file. The subsequent sorting process is thus activated by interpreting the chip ID, which ascertains the synthesis order of nucleotides in each microchip and directs them to the designated coupling reaction by automated transportation with a conveyor belt. Rigorous testing has validated the system’s exceptional accuracy, with a sorting error rate of less than 0.01%.

Step 3: Synthesis

Employing standard phosphoramidite chemistry within the mMPS synthesis module, a quintessential synthesis cycle encompasses four discrete steps: deblocking, coupling, oxidation, and capping. The procedure commences with the introduction of 3% trichloroacetic acid (TCA) in 2.5 mL of dichloromethane to the reaction column at ambient temperature, which is reacted for 30 seconds and reiterated twice. Subsequently, the TCA solution is evacuated, succeeded by a series of washes with 2 mL of acetonitrile (ACN), repeated five times, and the column is subjected to vacuum drying for 1 minute. Later, 0.7 mL of a 0.5 M activator solution in ACN is introduced and activated for a duration of 15 seconds. Post activation, 1.3 mL each of CAP A and CAP B are sequentially added to the reaction column for a 60-second interval, after which the solutions are drained. This is complemented by a washing procedure with 3 mL of ACN, repeated twice to ensure thorough cleansing. The coupling reaction is followed by the introduction of 2 mL of a 0.01 M oxidizing reagent, comprising a mixture of H_2_O, pyridine, and tetrahydrofuran in a volumetric ratio of 1:10:39, to the columns for a reaction period of 90 seconds. Subsequent to oxidation, the chips are rinsed with 2 mL of ACN, a process repeated five times to remove any residual reagents. Upon the completion of a base addition cycle, the chips are once again subjected to vacuum drying for a minute to prepare for subsequent cycles.

Step 4: Recycling

The recycling module, consisting of a mechanical arm and a vibratory bowl, plays a pivotal role in the post-synthesis phase, ensuring the efficient recycling of microchips and preparing for subsequent synthesis cycles. The mechanical arm adeptly transfers the chips from the reaction columns to the vibratory bowl which is designed with contours that precisely match the microchips’ geometries, thereby enhancing the system’s overall efficiency and minimizing the risk of misalignment or damage during the transfer process. Upon completion of these four modules, the pre-ordered nucleotides are successfully coupled to the chip surface.

### Oligo segmentation and synthesis

Oligo segmentation for optimal oligo mix concentration assay and high-throughput gene assembly were designed based on their respective sequences from GenBank (GFP: U17997.1, GH186: WP_000081498.1, RpoS: NP_415566.1, LacZ: U03993.1). The assembly oligonucleotides were designed using the GCATbio.Ltd customized software “gene2oligo”. For optimal oligo mix concentration assay, generated oligos were synthesized by GCATbio.Ltd and delivered at a concentration of 100 μM in nuclease-free water. For high-throughput gene assembly, oligos were synthesized using mMPS system.

### Optimal oligo mix concentration assay

A serial dilution of the oligonucleotide mix, ranging from 100 μM to 100 pM, was prepared to achieve final concentrations ranging from 1 μM to 1 pM per oligonucleotide. These dilutions were tested using the PCA-PCR protocol. The reaction products were analyzed by electrophoresis on 1% agarose gels stained with GelStain Blue (10,000×). A 2 μl aliquot of each reaction product was loaded onto the gel for analysis.

### PCA assembly

The primary PCA reaction was carried out in a total volume of 20 μL, containing a fixed concentration of 125 μM of each dNTP and 0.2 units of Q5 High-Fidelity DNA Polymerase (NEB), in 1x Q5 reaction buffer provided by the manufacturer. The thermocycling conditions were as follows: an initial hot start at 98 °C for 30 seconds, followed by denaturation at 98 °C for 2 seconds, annealing at 55 °C for 30 seconds, and extension at 72 °C for 30 to 90 seconds depending on gene length. A final extension was carried out at 72 °C for 5 minutes.

For the follow-up PCR, the reaction volume was 25 μL, which included 1 μL of the PCA reaction mixture as the template, and 0.25 units of Q5 High-Fidelity DNA Polymerase (NEB) in 1x Q5 reaction buffer. The thermocycling protocol consisted of an initial hot start at 98 °C for 30 seconds, denaturation at 98 °C for 10 seconds, annealing at 62 °C for 30 seconds, extension at 72 °C for 30 to 120 seconds depending on gene length, and a final extension at 72 °C for 5 minutes.

### High-throughput gene assembly

The gene set comprised two categories: 185 sequences of 600 – 1,200 bp and 100 sequences of 1800 – 3,200 bp. Gene assembly followed the PCA-PCR protocol. In the microarray method, oligonucleotides were sourced from Dynegene (ink-jet printing) and Genscript (electrochemistry), with design and amplification conducted as described previously. Plate subpools were amplified from 1 μL of the original template in 50 μL PCR reactions containing 0.4 units of Phusion High-Fidelity DNA Polymerase (NEB), 1x Phusion HF buffer, and 200 μM each dNTP. Assembly subpools were then amplified in 100 μL reactions using Phusion polymerase PCR. Restriction enzymes (BbsI-HF, BsmBI-v2, BsrDI, BspQI, BsaI-HFv2, and Nb.BtsI) (NEB) were used to selectively digest plate subpools, removing forward and reverse primer binding sites, followed by PCR to amplify the gene fragments.

### Mutagenesis library construction

The desired proteins are redesigned with NNK-type codon mutational scanning ranging from 57 to 297 bases in the CDS region, generating nine variant libraries including in total 66,691 sequences for de novo synthesis^38^. The nine libraries, originally labeled as A1, B3, and C1–C7, were named as Lib1 through Lib9 respectively. Additionally, a pair of 22-nt homologous sequences are introduced to the 5’ and 3’ end of each sequence in the libraries (GGGCTGCTCTAGAATGGCTAGC and AAGCTTGGCGGTGGCGGGTCTG respectively) to enable the next step ligation with expression vector pGJJ162. The generated libraries were in parallel synthesized using the developed mMPS system, in which both standard bases and degenerate bases N/K (representing the mixture of A, T, C and G, or the mixture of T and G respectively) were used for synthesis. The degenerate bases were pre-mixed in an optimized ratio to ensure equal representation of each nucleotide at the targeted mutation sites. Lib1, containing sequences shorter than 128 nt, was synthesized using single-stranded oligonucleotides directly cleaved from the microchip. In contrast, the remaining eight libraries (Lib2 to Lib9), composed of longer sequences (sequence lengths ranging from 179 to 341 bases), were divided into 378,896 sub-sequences, each ranging from 60 to 80 nt, for synthesis using the mMPS system. The full-length DNA fragments of each library were then assembled from these sub-fragments using PCA-PCR. For the other variant libraries, those ordered from IDT were designed as NNK-type libraries, while those from Twist Biosciences were created as pooled oligo libraries. The Twist libraries were designed to include all 19 possible amino acid substitutions at each position, utilizing the most abundant yeast codons for each substitution site^40^.

## Author contributions

Conceptualization: Y.S. J.W. and X.X.; Methodology: Y.S., X.Z., X.J. and Y.W.; Validation:Q.Z., R.Z., A.B., H.J., W.Y., N.C., H.Z., C.L., Y.W., C.X., K.Z., J.L., W.X., J.H., M.S., D.C., L.Z., S.M., N.G., and B.L.; Formal analysis: X.Z., X.J. and Y.W.; Data curation: X.Z., X.J., Y.W., and H.J.; Writing-original draft: X.Z., X.J. and Y.S.; Draft revision: All authors.

## Acknowledgments

We thank Ms. Z. Zhang and Ms. L. Yang for the efforts on schematic illustration design in Figure 1 panel A. This work was supported by National Key R&D Program of China (No. 2023YFF1206100), National Natural Science Foundation of China (No. 32322047 and 32401214), Shenzhen Science and Technology Program (Grant No.RCYX20210609103822039), and Jiangsu Provincial Department of Science and Technology (No.BM2023009).

## Reference

1. Jumper, J. et al. Highly accurate protein structure prediction with AlphaFold. Nature 596, 583–589 (2021).

2. Moger-Reischer, R.Z. et al. Evolution of a minimal cell. Nature 620, 122–127 (2023).

3. Fredens, J. et al. Total synthesis of Escherichia coli with a recoded genome. Nature 569, 514–518 (2019).

4. Schindler, D. et al. Design, construction, and functional characterization of a tRNA neochromosome in yeast. Cell 186, 5237-5253. e5222 (2023).

5. Taghon, G.J. & Strychalski, E.A. Rise of synthetic yeast: Charting courses to new applications. Cell Genom. 3 (2023).

6. Shen, Y. et al. Deep functional analysis of synII, a 770-kilobase synthetic yeast chromosome. Science 355, eaaf4791 (2017).

7. Ping, Z. et al. Towards practical and robust DNA-based data archiving using the yin–yang codec system. Nat. Comput. Sci. 2, 234–242 (2022).

8. Guarini, S., Bertolini, A., Lancellotti, N., Rompianesi, E. & Ferrari, W. Different cholinergic pathways are involved in the improvement induced by CCK-8 and by ACTH-(1–24) in massive acute hemorrhage, in rats. Pharmacol. Res. Commun. 19, 511–516 (1987).

9. Palluk, S. et al. De novo DNA synthesis using polymerase-nucleotide conjugates. Nat. Biotechnol. 36, 645–650 (2018).

10. McBride, L.J. & Caruthers, M.H. An investigation of several deoxynucleoside phosphoramidites useful for synthesizing deoxyoligonucleotides. Tetrahedron Lett. 24, 245–248 (1983).

11. Kosuri, S. & Church, G.M. Large-scale de novo DNA synthesis: Technologies and applications. Nat. Methods 11, 499–507 (2014).

12. Pease, A.C. et al. Light-generated oligonucleotide arrays for rapid DNA sequence analysis. Proc. Natl. Acad. Sci. 91, 5022–5026 (1994).

13. Singh-Gasson, S. et al. Maskless fabrication of light-directed oligonucleotide microarrays using a digital micromirror array. Nat. Biotechnol. 17, 974–978 (1999).

14. Hughes, T.R. et al. Expression profiling using microarrays fabricated by an ink-jet oligonucleotide synthesizer. Nature biotechnology 19, 342–347 (2001).

15. Ghindilis, A.L. et al. CombiMatrix oligonucleotide arrays: genotyping and gene expression assays employing electrochemical detection. Biosens. Bioelectron. 22, 1853–1860 (2007).

16. Tian, J. et al. Accurate multiplex gene synthesis from programmable DNA microchips. Nature 432, 1050–1054 (2004).

17. Kosuri, S. et al. Scalable gene synthesis by selective amplification of DNA pools from high-fidelity microchips. Nat. Biotechnol. 28, 1295–1299 (2010).

18. Quan, J. et al. Parallel on-chip gene synthesis and application to optimization of protein expression. Nat. Biotechnol. 29, 449–452 (2011).

19. Egeland, R.D. & Southern, E. Electrochemically directed synthesis of oligonucleotides for DNA microarray fabrication. Nucleic Acids Res. 33, e125–e125 (2005).

20. Moorcroft, M.J. et al. In Situ Oligonucleotide Synthesis on Poly(dimethylsiloxane): A Flexible Substrate for Microarray Fabrication. Nucleic Acids Res. 33, e75 (2005).

21. Sandahl, A.F. et al. On-demand synthesis of phosphoramidites. Nat. Commun. 12, 2760 (2021).

22. Fodor, S. et al. Light-directed, spatially addressable parallel chemical synthesis. Science 251, 767–773 (1991).

23. Fodor, S.P. et al. Multiplexed biochemical assays with biological chips. Nature 364, 555–556 (1993).

24. Roy, S., Gravener, L., Philp, D. & Kay, E.R. A dissipative reaction network drives transient solid–liquid and liquid–liquid phase cycling of nanoparticles. Angew. Chem. Int. Ed. 62, e202217613 (2023).

25. Qin, X., Vegge, T. & Hansen, H.A. Cation-Coordinated Inner-Sphere CO<sub>2 </sub>Electroreduction at Au–Water Interfaces. J. Am. Chem. Soc. 145, 1897–1905 (2023).

26. Pon, R.T. Solid-Phase Supports for Oligonucleotide Synthesis. Curr. Protoc. Nucleic Acid Chem. 53, 3–1 (2013).

27. She, X., Huang, A.Q., Lucia, O. & Ozpineci, B. Review of silicon carbide power devices and their applications. IEEE Trans. Ind. Electron. 64, 8193–8205 (2017).

28. Pienaar, E., Theron, M., Nelson, M. & Viljoen, H.J. A quantitative model of error accumulation during PCR amplification. Comput. Biol. Chem. 30, 102–111 (2006).

29. TerMaat, J.R., Pienaar, E., Whitney, S.E., Mamedov, T.G. & Subramanian, A. Gene synthesis by integrated polymerase chain assembly and PCR amplification using a high-speed thermocycler. J. Microbiol. Methods 79, 295–300 (2009).

30. Zhao, W. & Xiao, X. Complete genome sequence of Thermococcus eurythermalis A501, a conditional piezophilic hyperthermophilic archaeon with a wide temperature range, isolated from an oil-immersed deep-sea hydrothermal chimney on Guaymas Basin. J. Biotech. 193, 14–15 (2015).

31. Leng, H., Wang, Y., Zhao, W., Sievert, S.M. & Xiao, X. Identification of a deep-branching thermophilic clade sheds light on early bacterial evolution. Nat. Commun. 14, 4354 (2023).

32. Halper, S.M., Hossain, A. & Salis, H.M. Synthesis success calculator: predicting the rapid synthesis of DNA fragments with machine learning. ACS Synth. Biol. 9, 1563–1571 (2020).

33. Botstein, D. & Shortle, D. Strategies and applications of in vitro mutagenesis. Science 229, 1193–1201 (1985).

34. Lenug, D. A method for random mutagenesis of a defined DNA segment using a modified polymerase chain reaction. Technique. JMCMB 1, 11–15 (1989).

35. Reznikoff, W. Tn 5 transposition: a molecular tool for studying protein structure–function. Biochem. Soc. Trans. 34, 320–323 (2006).

36. Fowler, D.M. & Fields, S. Deep mutational scanning: a new style of protein science. Nat. Methods 11, 801–807 (2014).

37. Acevedo-Rocha, C.G., Ferla, M. & Reetz, M.T. Directed evolution of proteins based on mutational scanning. Protein Eng. Methods Protoc., 87–128 (2018).

38. Beltran, A., Jiang, X.e., Shen, Y. & Lehner, B. Site saturation mutagenesis of 500 human protein domains reveals the contribution of protein destabilization to genetic disease., Preprint at https://biorxiv.org/content/10.1101/2024.1104.1126.591310v591312 (2024).

39. Ding, G., Zheng, M., Xu, W. & Shen, W. Fabrication of controllable free-standing ultrathin porous alumina membranes. Nanotechnology 16, 1285 (2005).

40. Topolska, M., Beltran, A. & Lehner, B. Deep indel mutagenesis reveals the impact of insertions and deletions on protein stability and function. Preprint at https://biorxiv.org/content/10.1101/2023.1110.1106.561180v561181 (2023).

