## Supplementary file for "Revolutionizing large-scale DNA synthesis with microchip-based massive in parallel synthesis system"

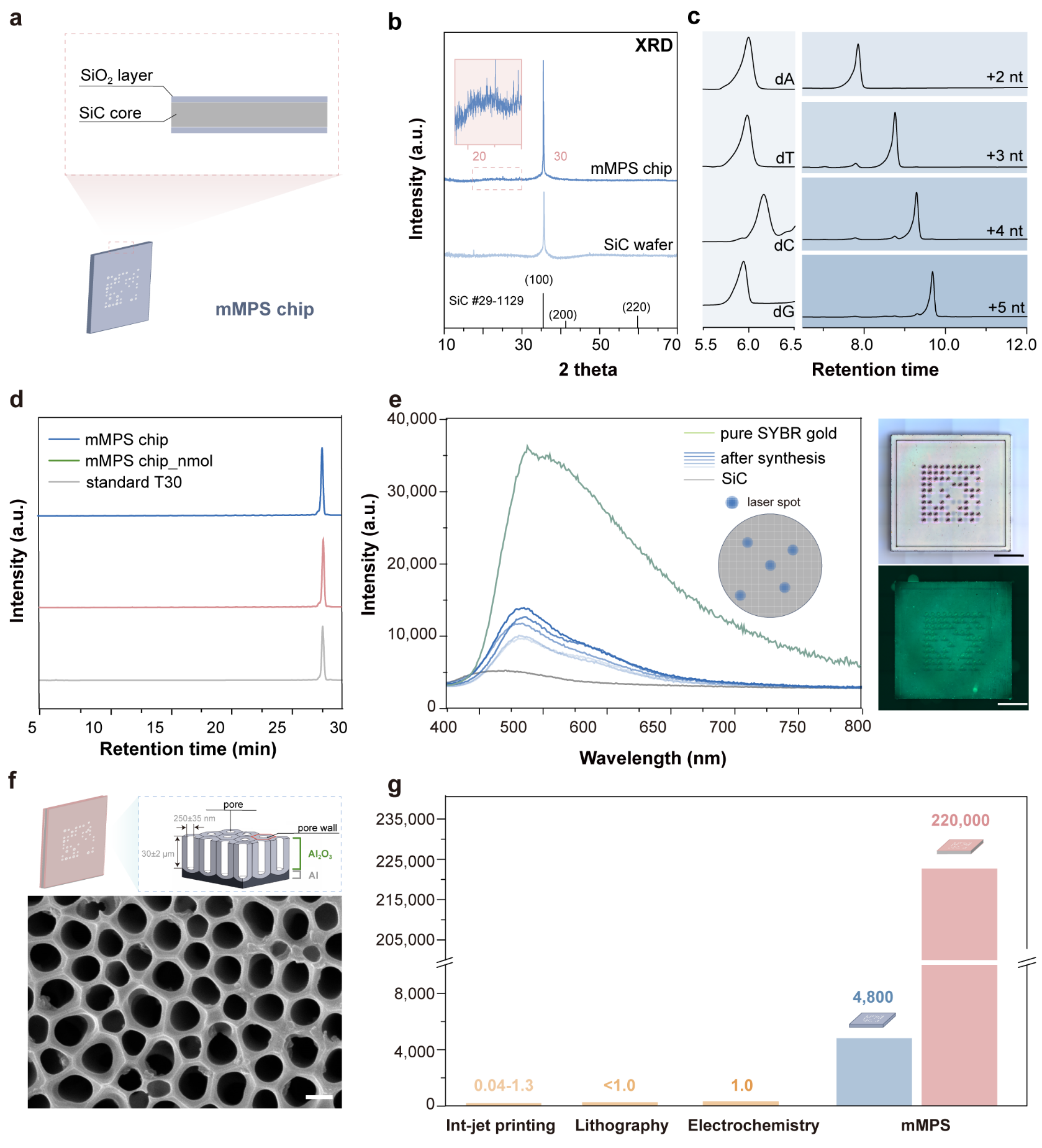

**Figure S1** (a) Schematics of the structural composition and (b) XRD pattern of mMPS chip.

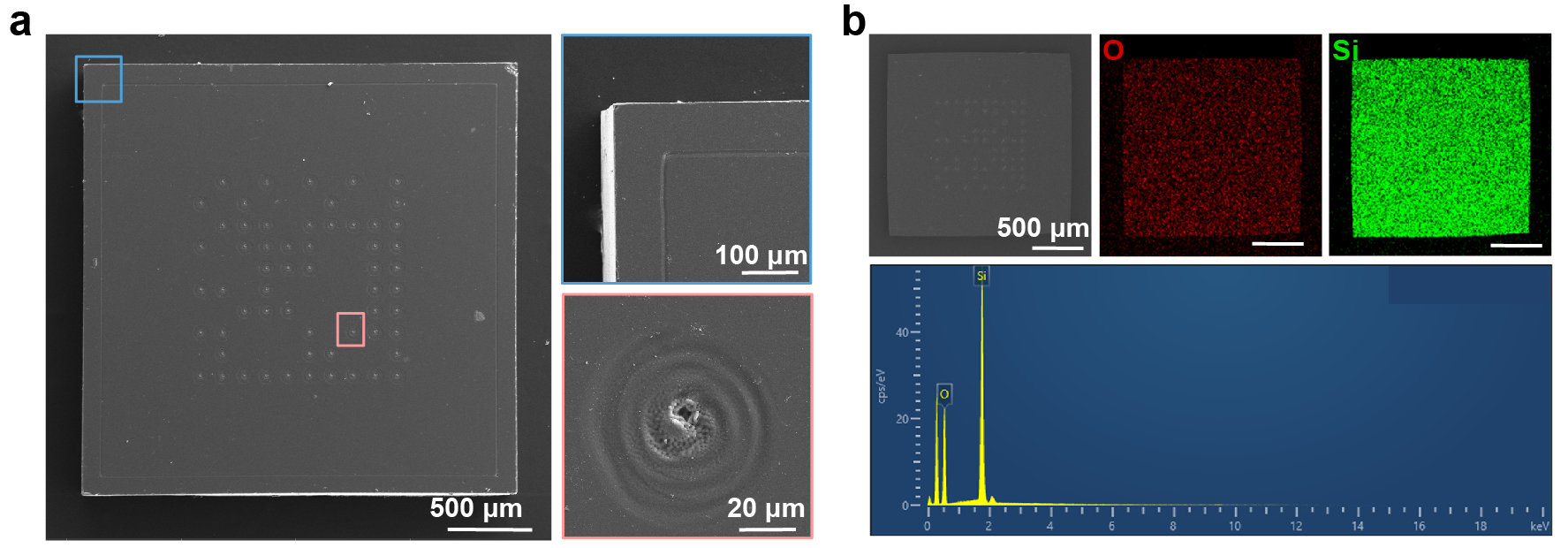

**Figure S2** (a) SEM images and (b) elemental maps of mMPS chip.

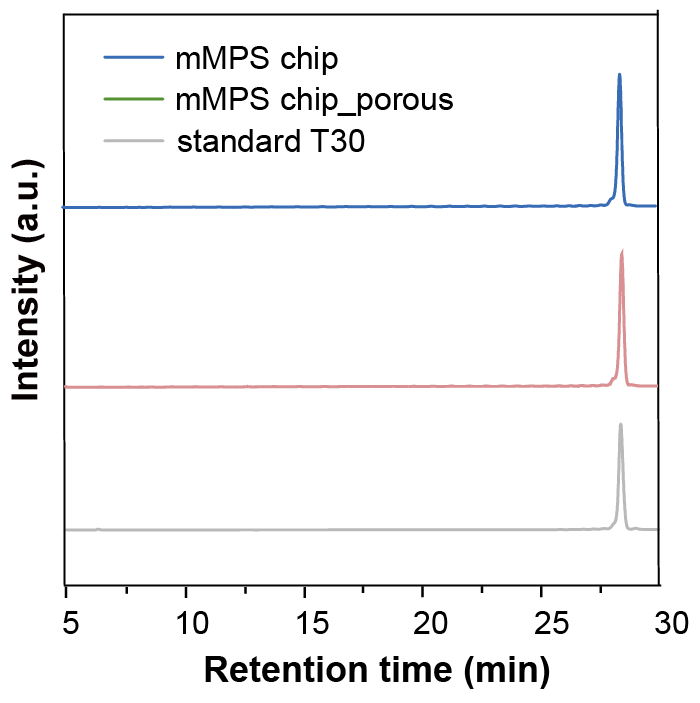

**Figure S3** HPLC spectra of T30 products synthesized by mMPS chip and mMPS_porous chip.

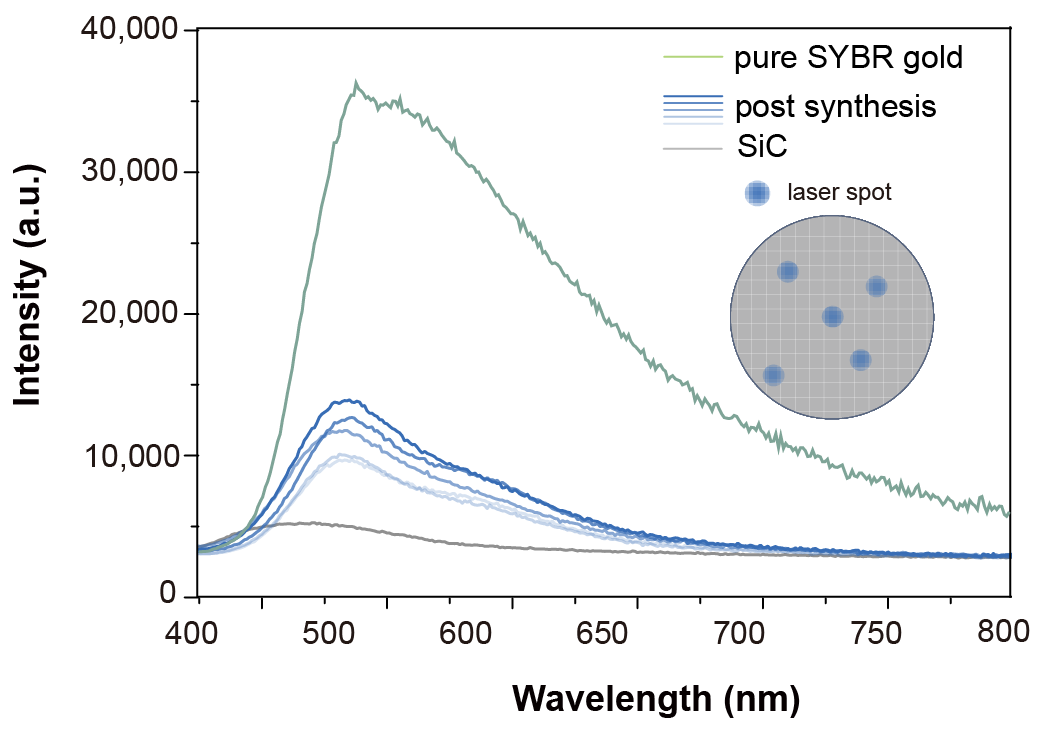

**Figure S4**. PL spectra of mMPS chip after oligo synthesis.

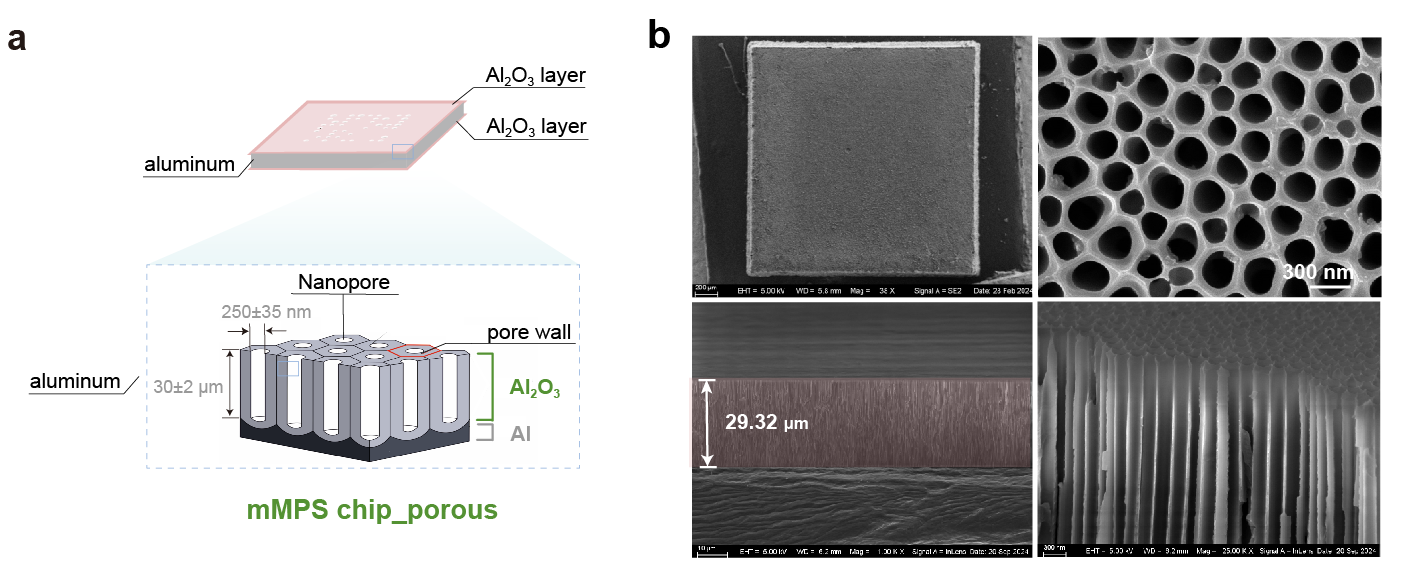

**Figure S5** (a) Schematics of structural composition and (b) SEM images of mMPS_porous chip on surface (top) and cross-section (down).

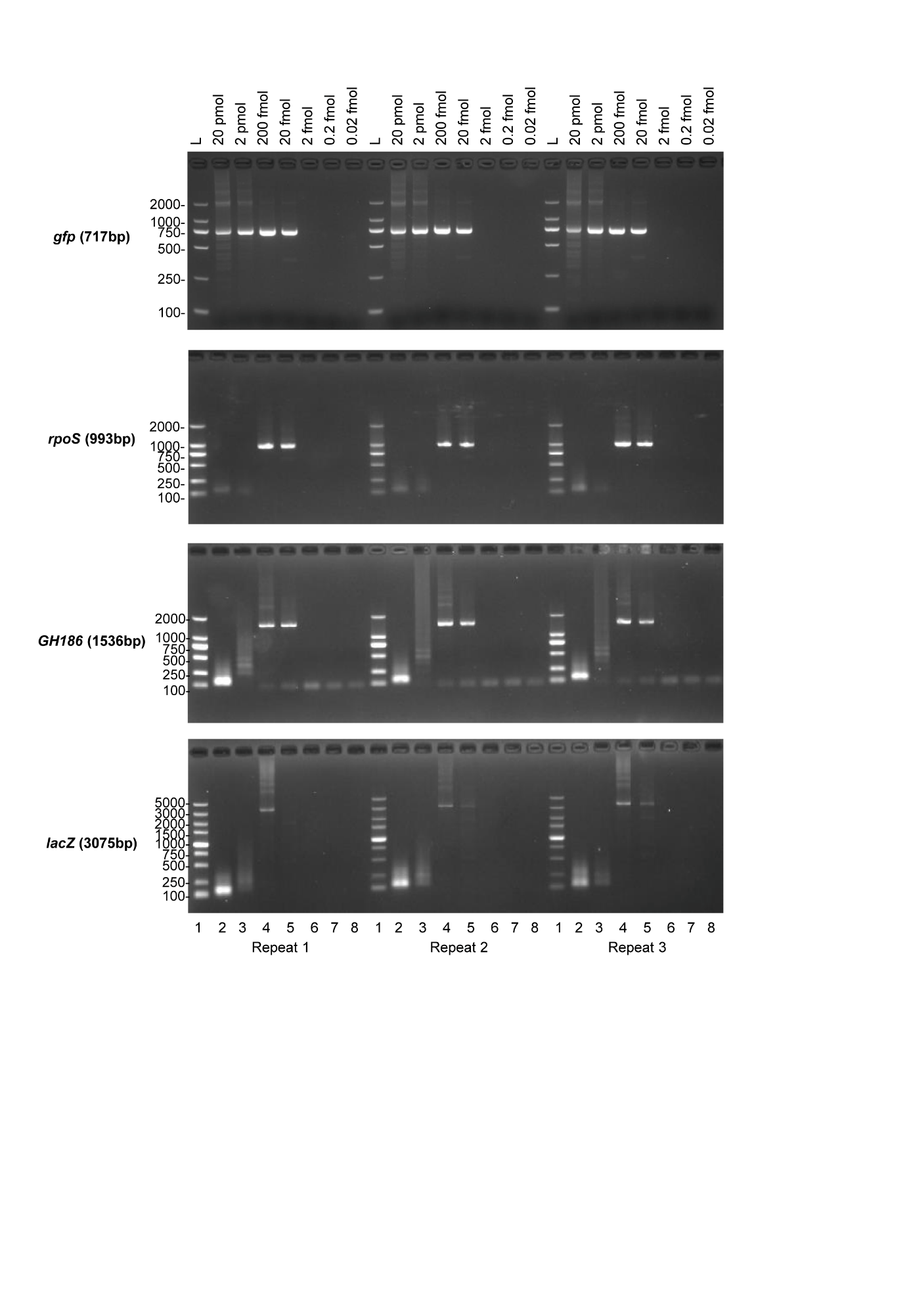

**Figure S6** Effect of oligo mix yield on gene assembly success rate. PCA-PCR products from four different genes - gfp (717 bp), rpoS (993 bp), GH186 (1536 bp), and lacZ (3075 bp) - were amplified using oligos with varying template concentrations. The volume of PCA reaction was set at 20 μL.

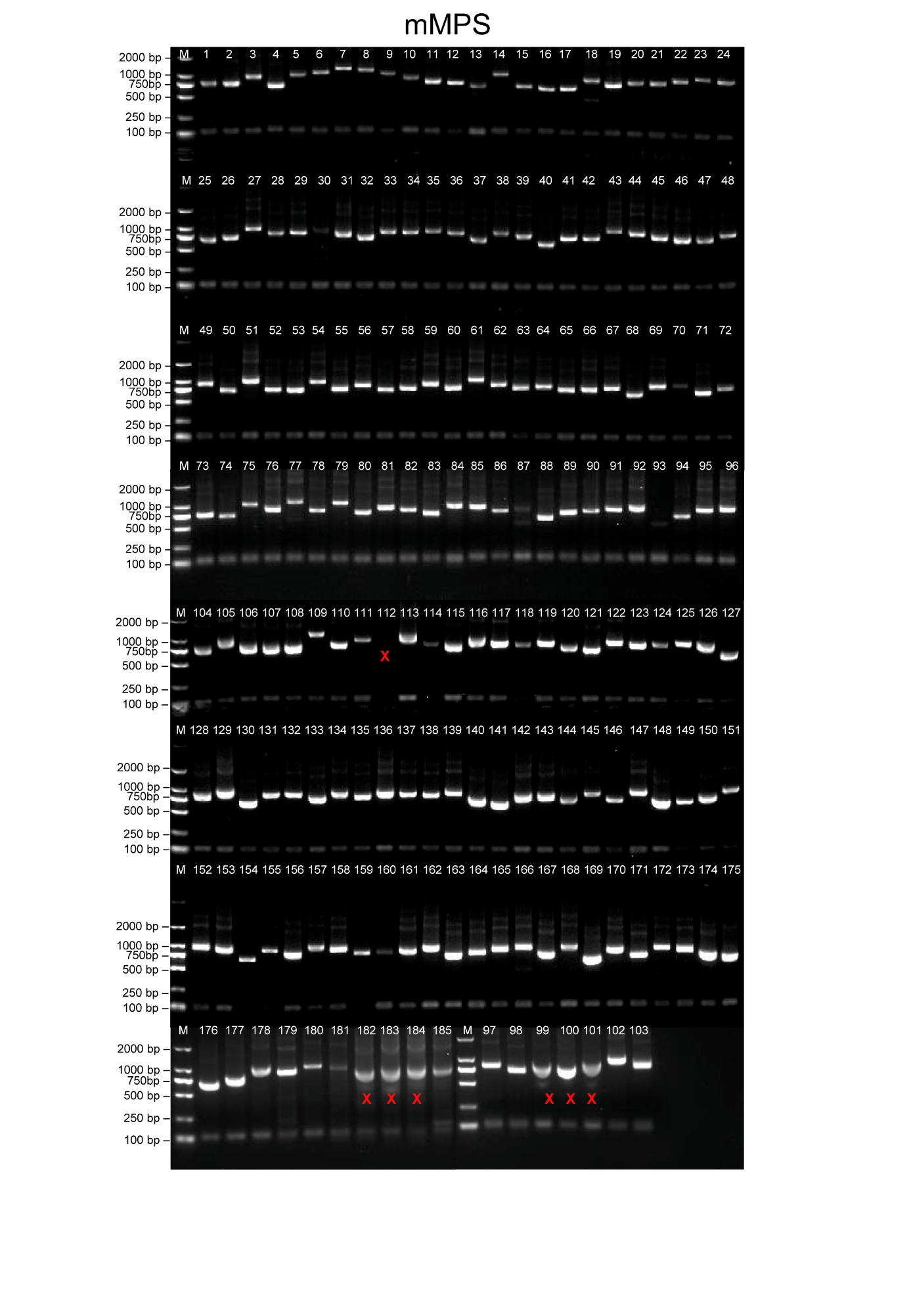

**Figure S7** Gene assembly using mMPS oligos. Assembly of 185 sequences, ranging from 600 to 1200 bp, was performed using the PCA protocol. Bands marked with an "X" indicate unsuccessful assembly outcomes.

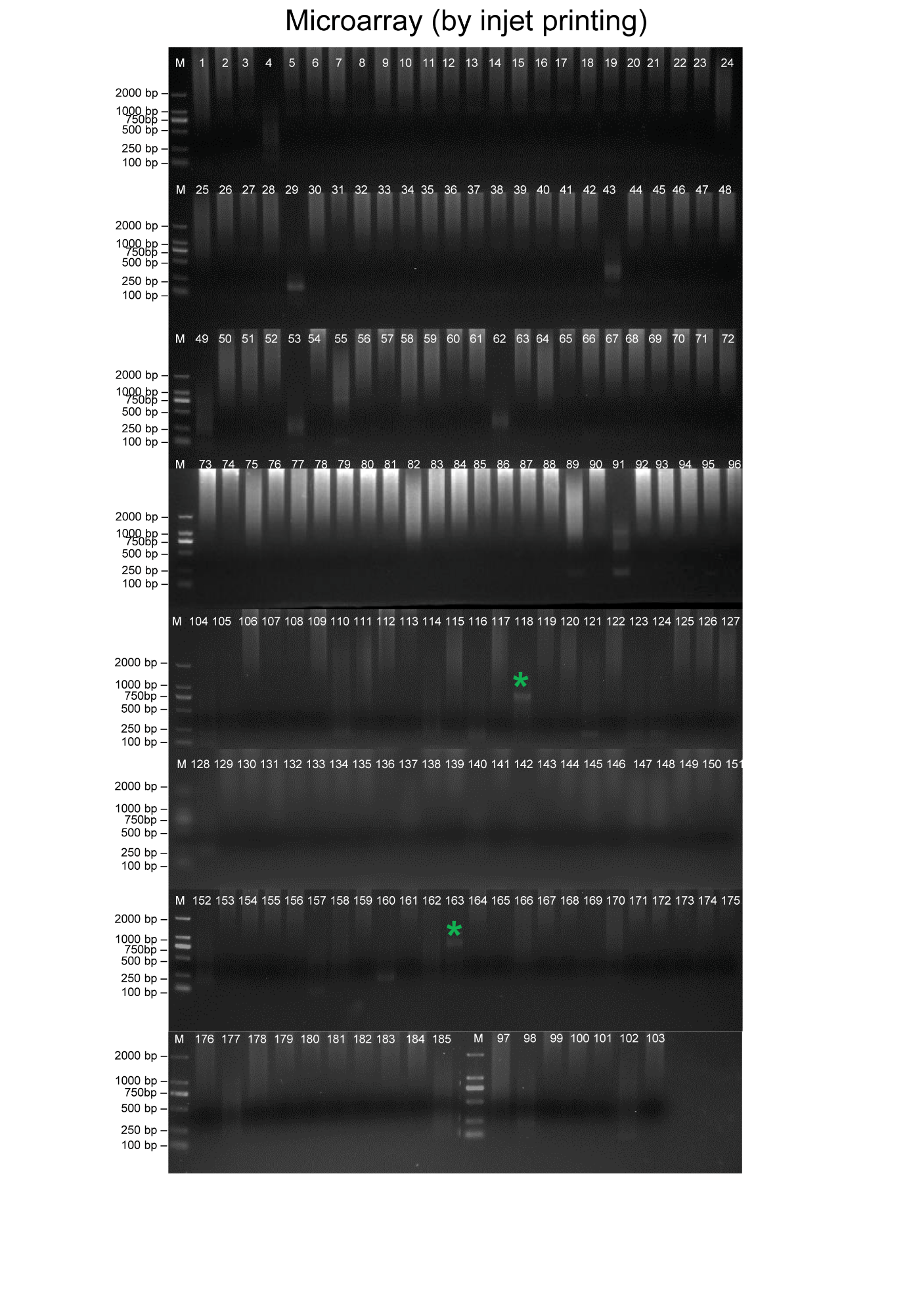

**Figure S8** Gene assembly using arrayed chip derived oligo by ink-jet printing approach. Assembly of 185 sequences, ranging from 600 to 1200 bp, was designed and performed as previously described^1^. Bands marked with an “*” denote successful assembly outcomes.

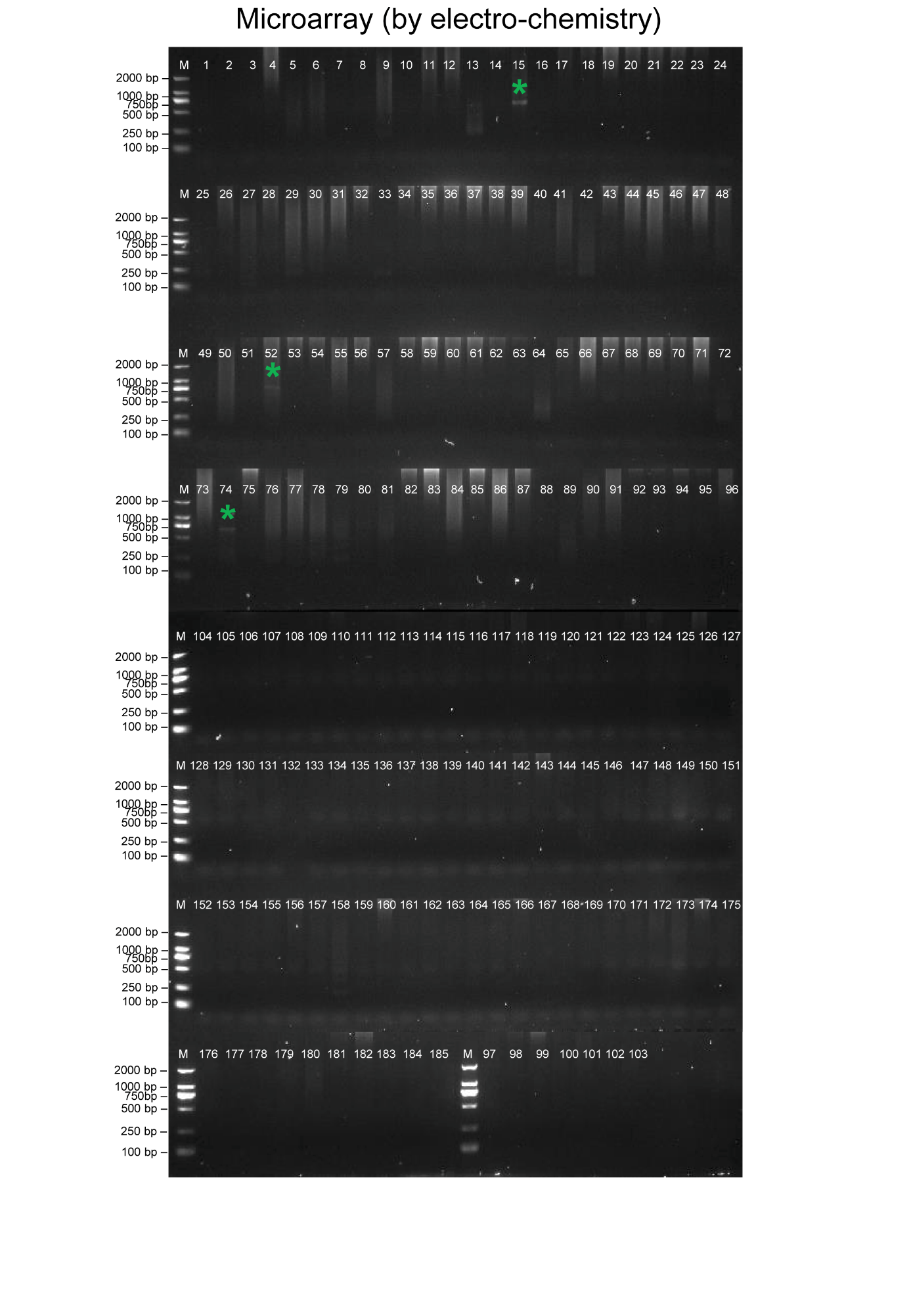

**Figure S9** Gene assembly using arrayed chip derived oligo by electrochemistry approach. Assembly of 185 sequences, ranging from 600 to 1200 bp, was designed and performed as previously described^1^. Bands marked with an “*” denote successful assembly outcomes.

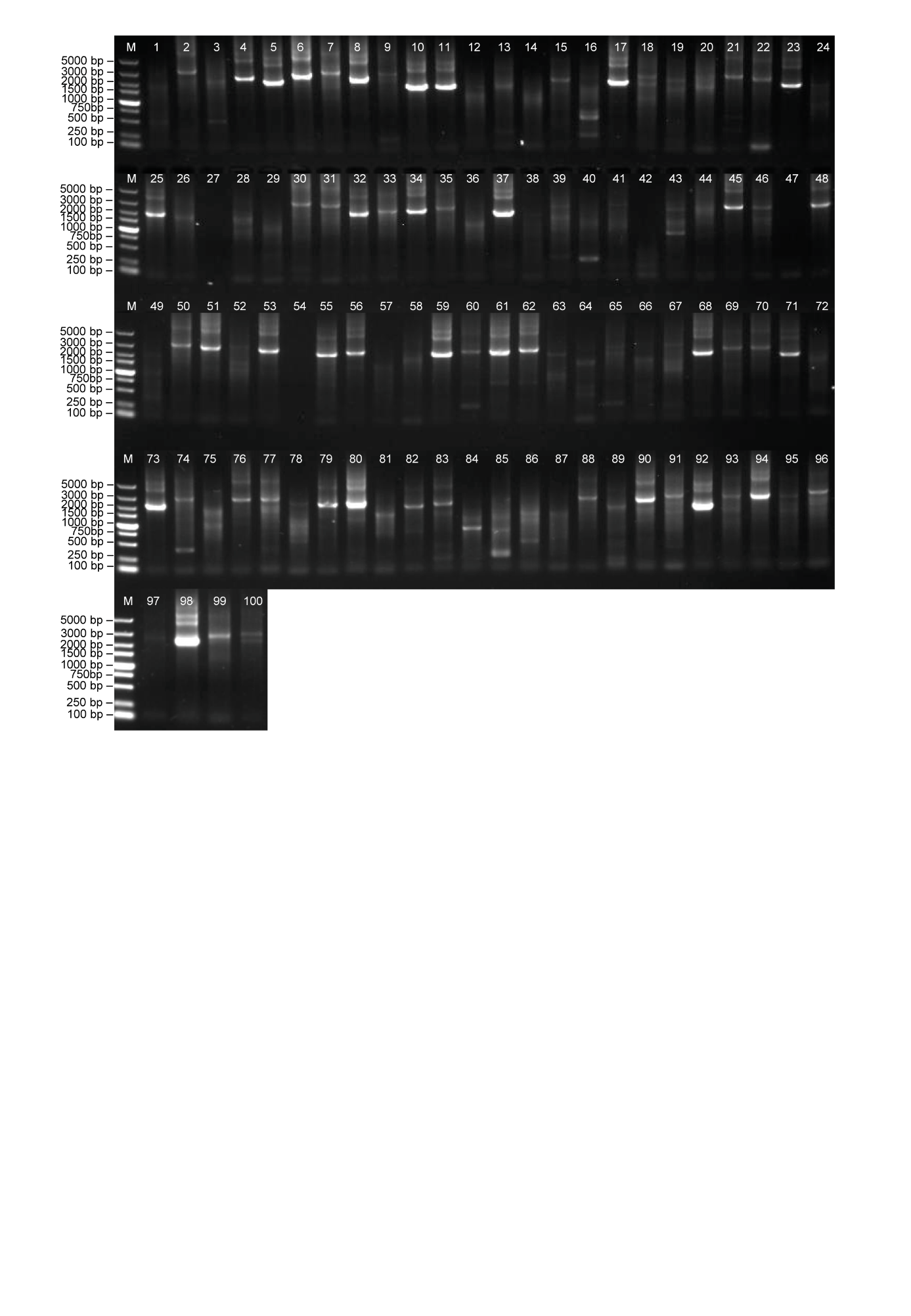

**Figure S10** Longer gene assembly using mMPS oligos. Assembly of 100 sequences, ranging from 1800 to 3200 bp, was performed using the PCA protocol.

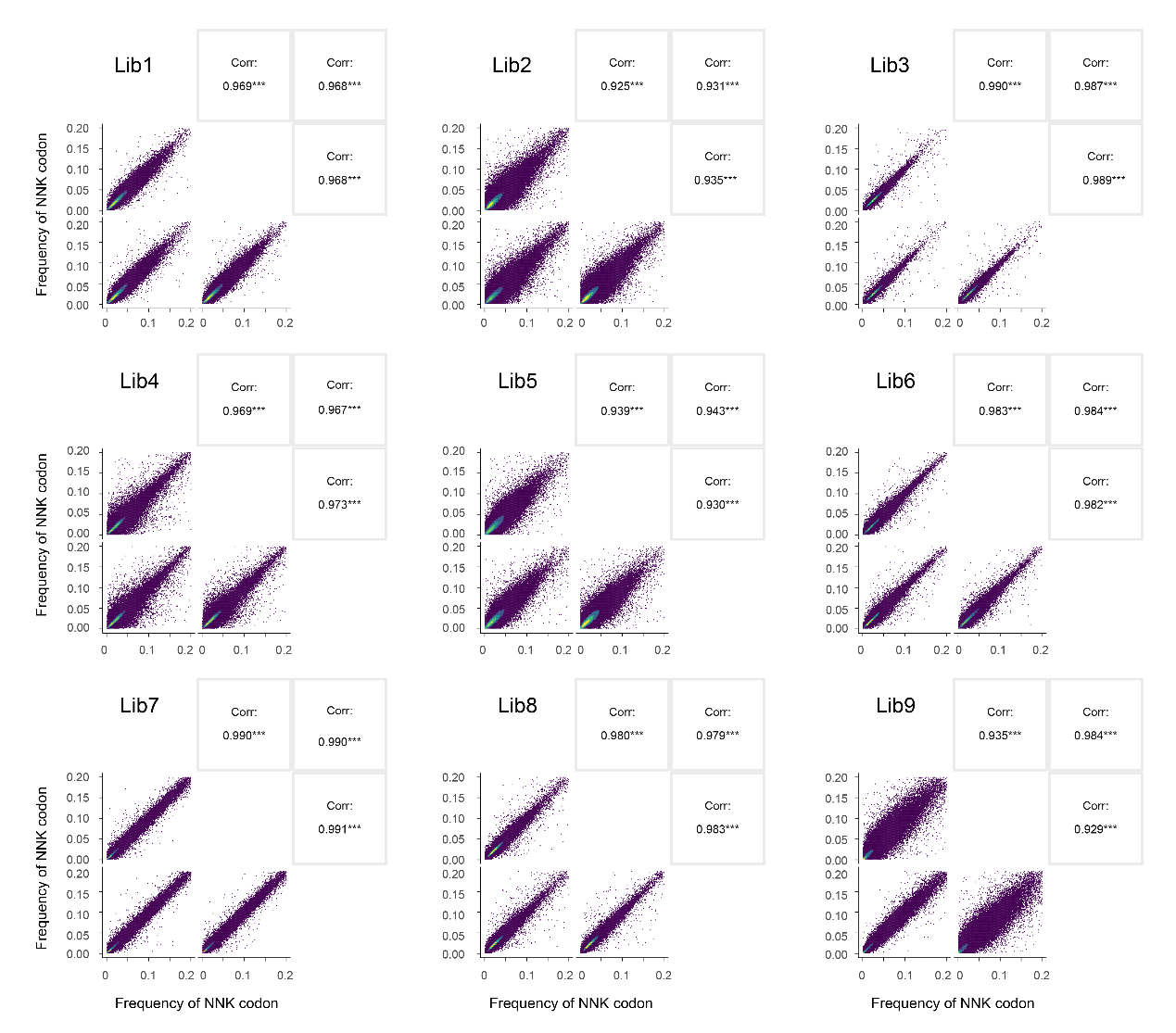

**Figure S11** Pearson correlation coefficients (r) between replicates for each of the nine libraries.

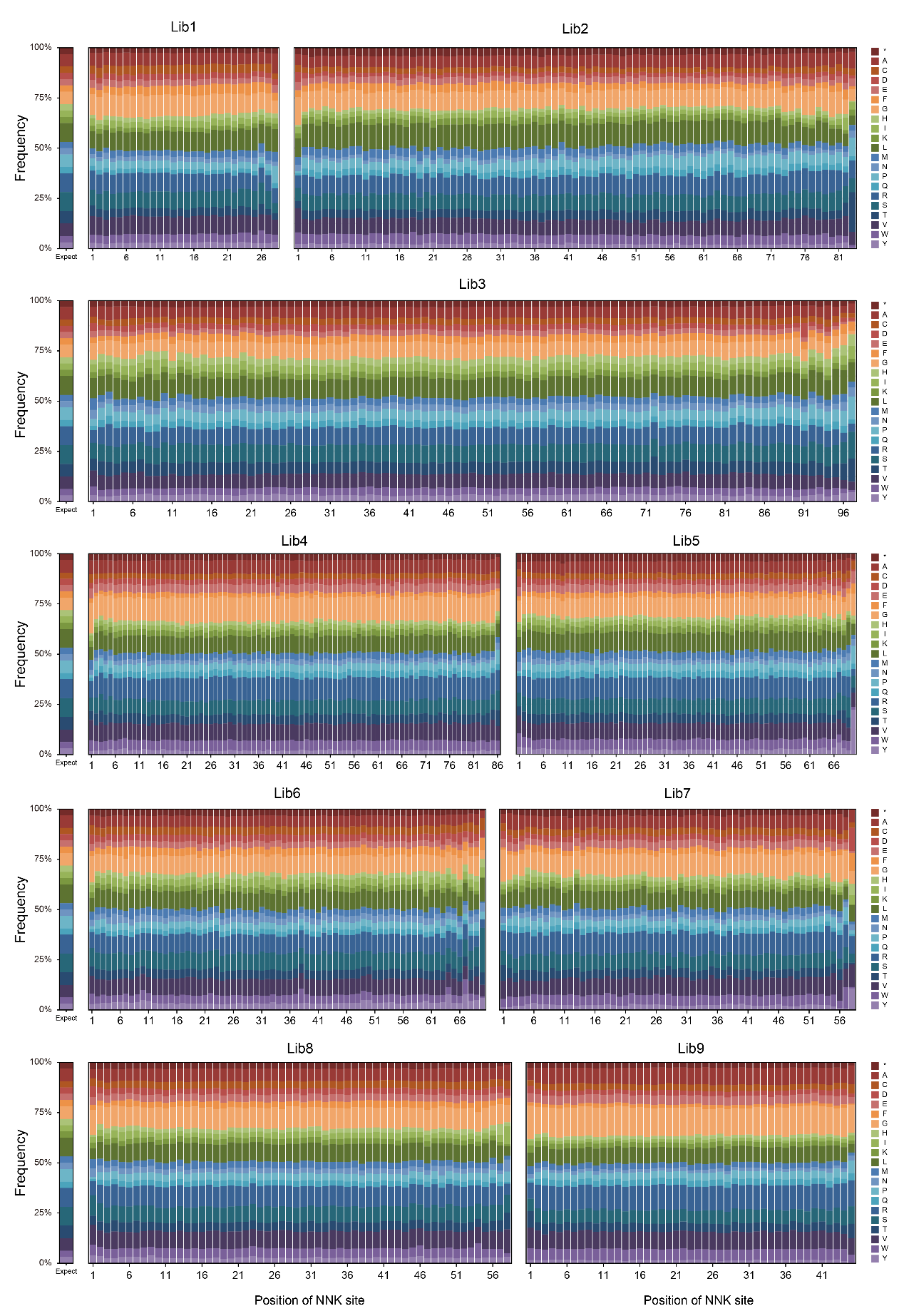

**Figure S12** Amino acid distribution analysis for each library, with theoretical expectations displayed on the left. The * symbol denotes stop codons.

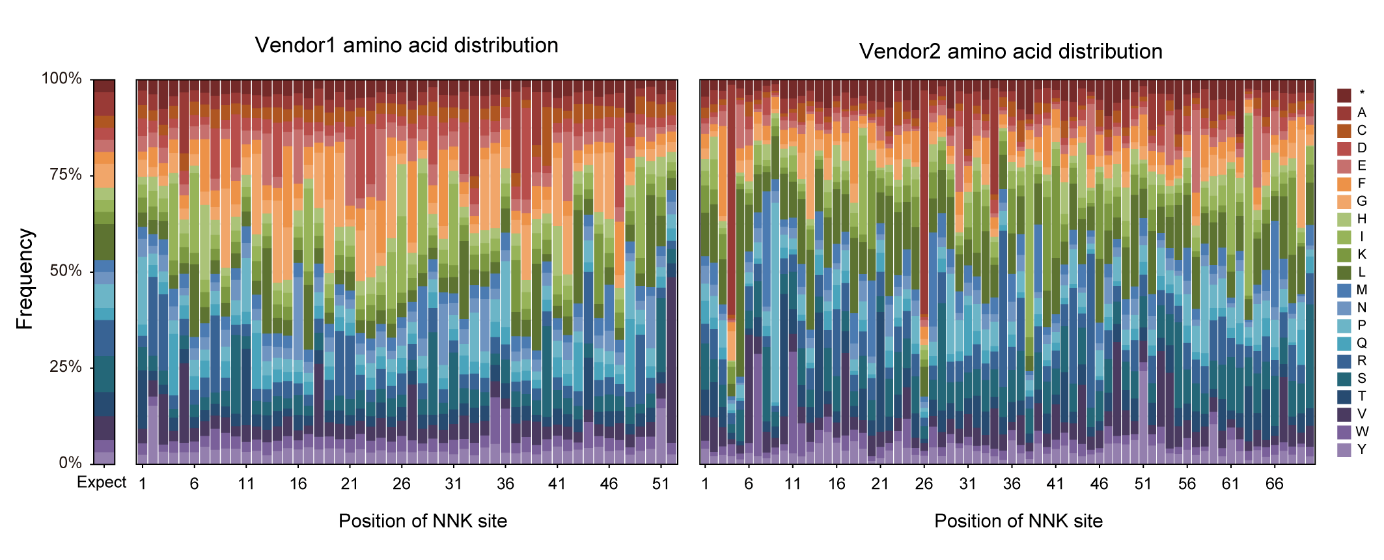

**Figure S13** Amino acid distribution analysis for variant library from two vendors using arrayed chip (vendor 1) and column based (vendor 2) DNA synthesis methods, with theoretical expectations displayed on the left. The * symbol denotes stop codons. Libraries from vendor 1 were designed to include all 19 possible amino acid substitutions at each position by specific oligos^2^. Libraries from vendor 2 were NNK-type libraries.

**Table S1** thickness measurements of mMPS chip through ellipsometry.

| **Measurement** | **Average thickness (nm)** |
| --- | --- |
| 1^st^ sample | 1043.6 |
| 2^nd^ sample | 1007.9 |
| 3^rd^ sample | 1013.2 |

The measurements were conducted on a random selection of three mMPS chips following the oxidation process to ensure a representative analysis of the sample set.

**Table S2**. Performance comparison of existing high-throughput DNA synthesis techniques.

| **Method** | **Yield/sequence (fmol)** |
| --- | --- |
| Ink-jet Printing^3^ | 0.04-1.3 |
| Lithography^4^ | ~5.0 |
| Electrochemistry^5^ | ~1.0 |
| **mMPS chip** | **1,216 ± 157** |
| **mMPS_porous chip** | **221,000 ± 415** |

**Reference**

1.	Kosuri, S. A scalable gene synthesis platform using high-fidelity DNA microchips. *Manuscript submitted for publication. Wyss Institute for Biologically Inspired Engineering*.

2.	Topolska, M., Beltran, A. & Lehner, B. Deep indel mutagenesis reveals the impact of insertions and deletions on protein stability and function. Preprint at <https://biorxiv.org/content/10.1101/2023.1110.1106.561180v561181> (2023).

3.	LeProust, E.M. et al. Synthesis of High-Quality Libraries of Long (150mer) Oligonucleotides by A Novel Depurination Controlled Process. *Nucleic Acids Res.* **38**, 2522-2540 (2010).

4.	Jia, J., Tong, C., Wang, B., Luo, L. & Jiang, J. Accurate Multiplex Gene Synthesis from Programmable DNA Microchips. *Nature* **432**, 1045-1050 (2004).

5.	Egeland, R.D. & Southern, E. Electrochemically directed synthesis of oligonucleotides for DNA microarray fabrication. *Nucleic Acids Res.* **33**, e125-e125 (2005).
